# Inactivation of posterior but not anterior dorsomedial caudate-putamen impedes learning with self-administered nicotine stimulus

**DOI:** 10.1101/2020.08.28.271908

**Authors:** Christopher L. Robison, Theodore Kazan, Rikki Miller, Nicole Cova, Sergios Charntikov

## Abstract

The rodent caudate-putamen is a large heterogeneous neural structure with distinct anatomical connections that differ in their control of learning processes. Previous research suggests that the anterior and posterior dorsomedial caudate-putamen (a- and p-dmCPu) differentially regulate associative learning with a non-contingent nicotine stimulus. The current study used bilateral NMDA-induced excitotoxic lesions to the a-dmCPu and p-dmCPu to determine the functional involvement of a-dmCPu and p-dmCPu in appetitive learning with contingent nicotine stimulus. Rats with a-dmCPu, p-dmCPu, or sham lesions were trained to lever-press for intravenous nicotine (0.03 mg/kg/inf) followed by access to sucrose 30 s later. After 1, 3, 9, and 20 nicotine-sucrose training sessions, appetitive learning in the form of a goal-tracking response was assessed using a non-contingent nicotine-alone test. All rats acquired nicotine self-administration and learned to retrieve sucrose from a receptacle at equal rates. However, rats with lesions to p-dmCPu demonstrated blunted learning of the nicotine-sucrose association. Our primary findings show that rats with lesions to p-dmCPu had a blunted goal-tracking response to a non-contingent nicotine administration after 20 consecutive days of nicotine-sucrose pairing. Our findings extend previous reports to a contingent model of nicotine self-administration and show that p-dmCPu is involved in associative learning with nicotine stimulus using a paradigm where rats voluntarily self-administer nicotine infusions that are paired with access to sucrose—a paradigm that closely resembles learning processes observed in humans.

## 1 Introduction

The use of tobacco products constitutes a major public health issue. The use of tobacco products is responsible for more deaths than alcohol and all illicit substances combined (Peacock et al., 2018). Globally, tobacco use is responsible for more than eight million deaths (WHO, 2019). Tobacco use is the third largest risk factor for cardiovascular disease worldwide (Benziger et al., 2016) and represents the largest cause of preventable death around the world (Lariscy, 2019). The compounding factor of the tobacco use problem is that many current tobacco users are trying to quit their debilitating habit but are unable to do so. For example, over 60% of current smokers report they desire to quit, yet, the rate of long-term success for each quit attempt is between 4%-7% (Courtney, 2015; Messer et al., 2008). Nicotine is the main constituent of tobacco products. Nicotine dependence is a multifaceted problem with genetic, biological, and societal aspects contributing to the perpetuation of tobacco use. Moreover, researchers are increasingly aware of the role of learning processes involving nicotine stimulus (Bevins et al., 2012), and there is ongoing effort to better understand neurobiological substrates underlying these learning processes (Charntikov et al., 2017, 2012). A better understanding of behavioral and neurobiological processes involved in learning with nicotine stimulus may eventually contribute to development of more efficacious treatment strategies.

Nicotine is a psychostimulant and a primary psychoactive and addictive constituent of tobacco products (Stolerman and Jarvis, 1995). Humans readily consume tobacco products in various forms, including smoking combustible products (e.g., cigarettes, cigars, hookah), chewing tobacco, inhaling vapors containing nicotine from electronically controlled heated devices, or inhaling aerosols from heat-not-burn products (e.g., heat sticks). Reinforcing effects of nicotine are often modeled in preclinical studies using the self-administration task. In these models, rats press one of the two available levers (i.e., active lever) to receive an intravenous infusion of a drug; the response on the other lever (i.e., inactive lever) usually has no programmed consequences. The reinforcing effects of a drug are evidenced by the higher responding on the active lever compared to an inactive lever (i.e., lever discrimination). Rats readily self-administer nicotine; however, the levels of nicotine self-administration are often lower than for other commonly abused substances. Indeed, extant research agrees that nicotine is a weak reinforcer when compared to other psychostimulants like cocaine or methamphetamine (Dougherty et al., 1981; Henningfield and Goldberg, 1983). In comparison to potent reinforcing effects of cocaine or methamphetamine, nicotine drives relatively low levels of behavioral output resulting in low levels of consumption (e.g., only 2-3 times higher than saline) and is often insensitive to the nicotine dose (Huynh et al., 2017; Kohut and Bergman, 2016). Nonetheless, there are some similarities between responding for nicotine and responding for other stimulants. For example, responding for nicotine is sensitive to schedules of reinforcement where the variable or progressive ratio of reinforcement schedules evoke higher levels of responding in comparison to fixed-ratio schedules (Charntikov et al., 2013; Donny et al., 2003, 1999; Kazan and Charntikov, 2019; Killeen et al., 2009). Although studying nicotine’s primary reinforcing properties and their behavioral and neurobiological effects is of great importance to understanding tobacco addiction, learning processes involving nicotine are likely to be more complex and may further explain the tenacity of the nicotine use habit. A better understanding of behavioral and neurobiological factors involved in learning with nicotine stimulus could improve existing models of nicotine use and allow for a more comprehensive assessment of novel treatment strategies.

In addition to mild reinforcing effects, nicotine enhances responding for other rewards and increases the incentive salience of other reinforcers (Donny et al., 2003; Garcia-Rivas and Deroche-Gamonet, 2019; Palmatier et al., 2007). For example, both contingent and non-contingent nicotine increase response rates for a visual stimulus without increasing overall locomotor activity (Constantin and Clarke, 2018; Donny et al., 2003). Furthermore, while prior research firmly established that nicotine stimulus can guide the behavior of rats in the form of goal-tracking or sign-tracking (Bevins et al., 2012; Dion et al., 2011; Palmatier et al., 2004), there is additional evidence that demonstrates nicotine’s ability to shift goal-tracking response to a sign-tracking response (Palmatier et al., 2013). Thus, pretreatment with nicotine can facilitate an approach to incentives associated with reinforcers, which indicates the increase in the incentive salience of stimuli that were previously associated with the reward. In addition to these reward or incentive enhancing effects, nicotine can function as an interoceptive stimulus that can acquire additional conditioned or reinforcing properties (Charntikov et al., 2020; Murray and Bevins, 2007). Early models investigating behavioral and neural mechanisms underlying learning with nicotine stimulus used a discriminated goal-tracking task. Using that task, rats would be pretreated with nicotine or saline prior to a training session where intermittent access to sucrose is only available on the day when rats receive nicotine. After a period of nicotine-sucrose pairings, rats would develop an anticipatory approach (i.e., goal-tracking) towards the area associated with a sucrose reward (i.e., sucrose receptacle). This behavioral response follows many postulates of Pavlovian learning and likely simulates learning processes in human tobacco users. Importantly, the neural substrates associated with this type of learning are essentially unexplored.

There is limited evidence suggesting that dmCPu is an integral part of a neurobiological mechanism involved in learning with nicotine stimulus. Our previous reports show that after a period of learning where interoceptive effects of non-contingent nicotine are paired with access to liquid sucrose, rats show higher nicotine-evoked neuronal activity in dmCPu (Charntikov et al., 2012). This involvement of dmCPu in associative learning with nicotine stimulus further strengthened previous accounts implicating dmCPu in instrumental stimulus-response and habitual learning (Everitt and Robbins, 2005; Ito et al., 2002, 2000). Assessment of functional involvement of dmCPu in learning with nicotine stimulus in a context of discriminated goal-tracking task shows that anterior (a-) and posterior (p-) dmCPu are differentially involved in the acquisition and expression of responding maintained by the nicotine stimulus (Charntikov et al., 2017). Specifically, the intact function of p-but not a-dmCPu is needed for the acquisition of learning with nicotine stimulus using non-contingent nicotine treatment in the context of a discriminated goal-tracking task. On the other hand, p- and a-dmCPu are differentially involved in the expression of learning with nicotine stimulus where transient lesions to p-dmCPu inhibit nicotine evoked responses while transient lesions to a-dmCPu increase goal-tracking rates on saline sessions by disinhibiting responding when nicotine stimulus is not detected (Charntikov et al., 2017). These findings, using the discriminated goal-tracking task used in previous reports, provided an important initial step towards an understanding of behavioral, pharmacological, and neural mechanisms involved in learning with nicotine stimulus. One of the limitations in using the discriminated goal-tracking task is a lack of choice to respond for a reinforcer. Providing a choice to respond for a reinforcer may better model processes in the clinical population and thus may engage more relevant behavioral and neural mechanisms in preclinical models of substance use. For example, smokers have a choice to engage in smoking behavior or not. Furthermore, smokers often experience nicotine state in the presence of abundantly available rewarding stimuli like food, alcohol, work breaks, or peer interaction, to name a few. To address these limitations, the existing nicotine self-administration task. In this model, rats self-administer nicotine, where each nicotine infusion is paired with access to liquid sucrose, and each nicotine infusion is temporally spaced out to increase the salience of each successive infusion. This approach allows us to study behavioral and neural mechanisms underlying learning with nicotine stimulus in a model that better parallels nicotine use in human smokers. Using this preclinical approach to study appetitive learning with nicotine stimulus, we hypothesized that a- and p-dmCPu would be differentially involved in acquisition of learning with nicotine stimulus. We tested this hypothesis by lesioning a- and p-dmCPu prior to the conditioning phase of the study where self-administered nicotine is paired with access to sucrose. We then assessed behavioral responses across a range of dependent variables and over the temporal progression of learning with nicotine stimulus.

## 2 Materials and Methods

### 2.1 Animals

Fifty-seven male Sprague Dawley rats (250-300 g) were purchased from Envigo (Indianapolis, IN, USA). Rats were single-housed in a temperature-controlled vivarium on a 12 h light/dark cycle (lights on at 0700). Rats were acclimated to the colony for one week before experimental procedures. Food and water were available ad libitum during the acclimation period and for one week after each surgery. Rats were water-deprived for 22 hours per day during lever training; water was available for one hour during lever training, and for one hour immediately after. Throughout the nicotine self-administration phase, rats were food-restricted to maintain 85 % of their free-feeding weight with water available ad libitum. Free-feeding weight was increased by 2 g every 30 days. All procedures were carried out in accordance with the Guide for the Care and Use of Laboratory Animals (National Research Council et al., 2010) under review and approval of the University of New Hampshire Institutional Animal Care and Use Committee.

### 2.2 Apparatus

Behavioral tests were carried out in Med Associates conditioning chambers measuring 30.5 × 24.1 × 21.0 cm (l × w × h) that were enclosed in a sound- and light-attenuated cubicle equipped with an exhaust fan (ENV-018MD; Med Associates, Inc.; St. Albans, VT, USA). Each chamber had aluminum sidewalls, metal rod floors, and polycarbonate for all other surfaces. A recessed receptacle (5.2 × 5.2 × 3.8 cm; l × w × d) on one wall provided access to a dipper arm. When raised, the dipper arm provided access to 100 μL of tap water (lever training sessions) or 30 % (w/v) sucrose solution (self-administration sessions). Access to the dipper area was monitored via an infrared beam mounted 1.2 cm into the receptacle and 3 cm above the floor. Two retractable levers (147 nN required for micro-switch closure) were mounted on each side of the receptacle. Levers were used as manipulanda to operate the dipper (training trials) or infusion pump (self-administration trials). A house light (two white 28V, 100 mA lamps) was located 10 cm above the conditioning chamber ceiling. An infusion pump (PMH-100VS; Med Associates) fitted with a 5-mL syringe was located outside each enclosure. The infusion pump (PMH-100VS; Med Associates; St. Albans, VT, USA) for each chamber was located outside the sound-attenuating cubicle. A 5 mL syringe was connected to a swivel using Tygon* tubing (AAQ04103; VWR; West Chester, PA, USA). Each swivel was coupled with a spring leash (C313C; Plastics One; Roanoke, VA, USA), which was suspended over the ceiling of the chamber on a balanced metal arm. Med Associates interface and software (Med-PC for Windows, version IV) were used to collect data and present programmed events.

### 2.3 Drugs

Nicotine bitartrate (MP Biomedicals; Solon, OH, USA) was dissolved in 0.9 % sterile saline. The pH of nicotine was adjusted to 7.0 ± 0.2 with a dilute NaOH solution. Nicotine doses are reported as a base. Doses and administration protocols were adopted from previous research (Charntikov et al., 2014; Kazan and Charntikov, 2019; Liu et al., 2008).

### 2.4 Excitotoxic lesion surgery

Rats were randomly assigned to one of three surgical conditions: sham, a-dmCPu, and p-dmCPu lesions. Half of the rats in a sham condition received sham injections to a- and the other half to p-dmCPu. Anesthesia was induced with 5 % isoflurane for 5 minutes and maintained at 2.5 % for the remainder of the surgery. Butorphanol (5 mg/kg; SC) and meloxicam (0.15 mg/kg; SC) were administered after induction to manage pain. Rats were then mounted on a stereotaxic apparatus, and bilateral craniotomies were performed under aseptic conditions with placement calculations relative to the bregma suture and the skull surface. Rats then received bilateral injections of sterile water or NMDA (0.5 μL/side; 0.12 M NMDA in sterile water; 0.05 μL/min infusion rate) to either the anterior (β +1.2 mm A/P, ±1.9 mm M/L, −4.7 mm D/V) or posterior dmCPu (β −0.35 mm A/P, ±2.4 mm M/L, −4.6 mm D/V). The injection needle was left in place for an additional 2 min and then withdrawn over the final 2 min period. All injections were controlled by a dual syringe pump (KDS-200; KD Scientific, Holliston, MA, USA) and were confirmed via displacement of a small bubble that was created in the tubing leading to the injection needle (28 gauge). After the surgery and for the following two days, rats were treated once a day with butorphanol (5 mg/kg; SC). After the surgery, rats were monitored daily, and given at least one week to recover before additional lever training. Four rats were removed from the study because they were unable to recover from surgery.

### 2.5 Preliminary lever training

Rats were first water-deprived for 22 hours a day and then were trained to retrieve water from a dipper receptacle in a water-deprived state. Rats were trained to retrieve water from a dipper receptacle until reaching 80 % retrieval criterion (3-5 days). These dipper training sessions consisted of 50 min trials with non-contingent water presentations delivered on a variable time interval (~ 3 rewards per minute). Water was returned to home cages immediately after the session for the following 70 min. Rats were then trained to lever press for liquid sucrose (5 % w/v; 100 μL) using a lever shaping procedure. At the start of each session, the house-light was turned on, and a randomly selected lever (right or left) was inserted. A lever press or lapse of 15 s resulted in sucrose delivery (4-s access), lever retraction, and commencement of a timeout (average=60 s; range=30 to 89 s). Following the timeout, a randomly selected lever was inserted with the condition that the same lever could not be presented more than twice in a row, and the number of left and right lever presentations was equal across the session. This protocol was repeated for 60 sucrose deliveries. Sessions lasted 65 to 80 min, depending on individual performance. Training continued until rats made lever presses on at least 80 % of lever insertions for two consecutive days (total training time was 3 to 6 daily sessions based on individual performance). Three rats were removed from the study due to the inability to acquire lever-pressing behavior.

### 2.6 Catheter implantation surgery

Upon completion of lever training, rats were equipped with intravenous (IV) jugular catheters. Isoflurane anesthesia was used as described above. A polyurethane catheter with a rounded tip and dual suture beads (RJVR-23; Strategic Applications Inc.; Lake Villa, IL, USA) was implanted into the right external jugular vein. The catheter was routed around the ipsilateral shoulder and affixed to a polycarbonate access port (313-000B; Plastics One Inc.; Roanoke, VA, USA) implanted along the dorsal midline 1 cm posterior to the scapulae. Immediately following surgery, catheters were flushed with 0.2 ml cefazolin (50 mg/ml) that was diluted in sterile saline with heparin (30 U/ml). This catheter flushing protocol was performed daily to maintain catheter patency throughout the self-administration phase. Pain management was identical to the protocol used after the lesion surgeries. At the end of the self-administration phase, catheter patency was assessed using an infusion of 0.05 mL xylazine (20 mg/mL) infused through the IV catheter. This xylazine dose produces rapid and transient motor ataxia in rats with patent catheters (Kazan and Charntikov, 2019; Stafford et al., 2019). Four rats were excluded from the study due to the loss of catheter patency throughout the study.

### 2.7 Nicotine self-administration: conditioning phase

Rats self-administered nicotine (0.03 mg/kg/inf; 2 h sessions) over 21 daily sessions. Active levers were assigned pseudo-randomly to each rat to ensure equal numbers of left and right active levers in each lesion condition. Both levers were inserted into the chamber at the beginning of each session. Rats were allowed to selfadminister nicotine using a variable ratio (VR3) schedule of reinforcement (i.e., on average, every third response was followed by access to water; range=1 to 5 presses). Meeting a schedule requirement resulted in the withdrawal of the levers, turning on a house light for 3 sec, and a ~1-second infusion of nicotine. All rats selfadministered the exact dose of nicotine using a variation in infusion duration that was automatically calculated by the program based on their pre-session weight. Thirty seconds after each infusion of nicotine, a dipper arm containing 100 μL of liquid sucrose (30 % w/v) was raised for 10 seconds. During the 10-second sucrose presentation, any entry into the dipper area greater than 0.02 seconds was considered to be the retrieval of sucrose. Levers were inserted back into a chamber after 8 min timeout. This prolonged timeout was instituted to ensure the salience of each successive infusion with the goal of strengthening the nicotine-sucrose association (Charntikov et al., 2020; Murray and Bevins, 2009). The number of infusions was limited to ten. Immediately after each self-administration session, catheters were flushed with the cefazolin (10 mg) diluted in 0.2 mL of heparinized saline (30 U/mL).

### 2.8 Non-contingent nicotine-alone tests

Immediately prior to sessions 2, 4, 10, and 21 of nicotine self-administration, rats were subjected to 8-min non-contingent nicotine-alone tests. During these tests, entries into the sucrose receptacle in the absence of cues or levers previously used in the study were recorded. Four minutes after the beginning of the test, rats received a single non-contingent and unsignaled infusion of nicotine (0.03 mg/kg; IV). The responses were recorded for another four minutes in the absence of any cues. Immediately after this 8 min test, rats progressed to self-administering nicotine paired with access to sucrose as described earlier.

### 2.9 Histological assessment

One day after the completion of all experimental procedures, rats were euthanized with Euthasol (Virbac, Fort Worth, TX, USA) and perfused with 100 mL 0.02 M sodium phosphate-buffered saline (PBS; 0.9 % NaCl, pH 7.4) followed by 300 mL 4 % paraformaldehyde solution dissolved in 0.02 M PBS (pH 7.4). Immediately after perfusion, brains were collected and post-fixed in 4 % paraformaldehyde for 24 hours at 4° C, followed by cryoprotection in 30 % sucrose solution diluted in 0.02 M PBS and stored at 4° C until saturated. 40 μm coronal brain sections of the CPu were taken from β +2.5 mm, and β −1.5 mm and a subset of 8 sections from each brain (4 sections anterior to and 4 sections posterior to the cannula insertion site, 160 μm between serial sections) were selected for histological assessment. Sections were stained for NeuN immunohistochemistry using 1:10,000 polyclonal rabbit anti-NeuN antibody (ABN78; EMD Millipore; Temecula, CA, USA) and horse anti-rabbit secondary antibody via an HRP reagent (ImmPRESS MP-7401; Vector Laboratories, Burlingame, CA, USA). Positive labeling was then visualized using peroxidase-activated substrate (ImmPACT VIP SK-4605; Vector Laboratories). Stained sections were mounted to slides, coverslipped, and then photomicrographed using a Nikon A1R confocal microscope using green laser (488 nm) transmission at 100x magnification. Lesions were considered to meet a criterion when the area without visible NeuN staining was located inside the predetermined dmCPu quadrant (Figure 1 right panels) and when a lesion epicenter was identified to be between β +2.04 mm and β +0.72 mm (a-dmCPu) or β +0.48 mm and β −0.48 mm (p-dmCPu). Twelve rats were removed from the study because their lesion placement did not meet that criterion.

**Figure 1.**
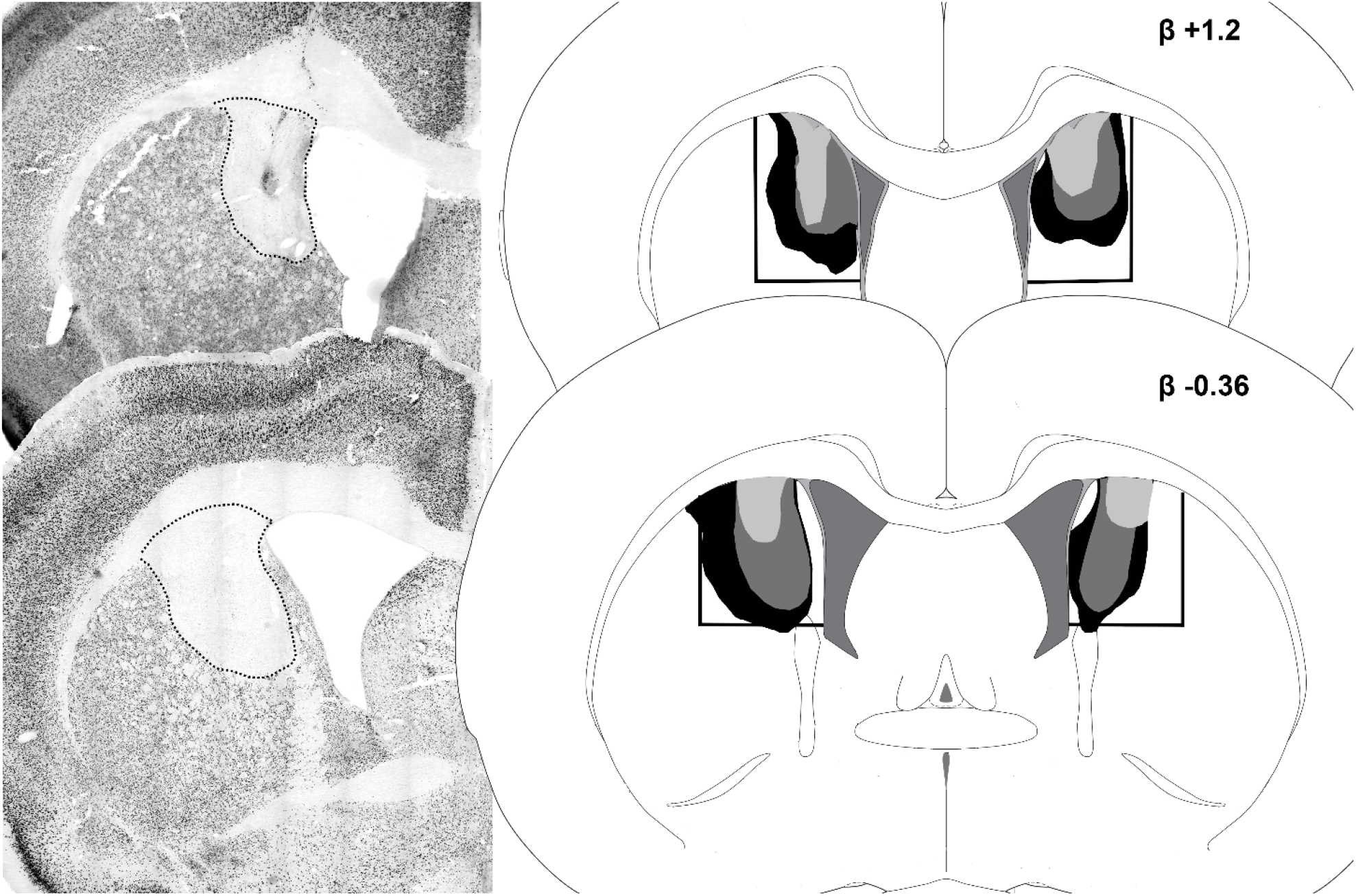
Photomicrographs of the representative NeuN stained a-dmCPu (top left) and p-dmCPu (bottom left) lesions and graphical illustrations of the extent of corresponding lesions (corresponding right panels). Dashed lines on the left panels show lesion outline and black lines on graphical illustrations (right panels) show targeted areas. Black filled areas on the right panels represent the largest extend of the damage, and grey filled areas represent smaller lesion sites observed in this study. Numbers on the right panels indicate targeted Bregma position.

### 2.10 Dependent Measures

The lesion size for each rat was measured in the photomicrograph pixel area at the coronal plane of the greatest lesion extent. During the conditioning phase where rats self-administered nicotine coupled with access to sucrose, the primary dependent measures were the number of lever presses on the active and inactive levers, the number of nicotine infusions, the number of sucrose retrievals, the duration of time spent in the dipper entry area, and the latency to reach the dipper entry area after each nicotine infusion. During non-contingent nicotine-alone tests, the total duration of head entry into the dipper receptacle was recorded by 15 s time bins numbered 1-32. The duration of head entries into the dipper receptacle provides a measure of goal-tracking activity. In comparison to the total number of dipper entries or dipper entry frequency, the duration of head entries into the dipper receptacle provides a more detailed account of behavioral response because the number of dipper entries measure does not account for the total time spent in the receptacle. For example, a rat may have one dipper entry after a nicotine infusion but may spend all the time between the nicotine infusion and a sucrose presentation with the snout in the receptacle, demonstrating a goal-tracking response. For this reason, we used a duration of head entries into the dipper receptacle as a main dependent measure of goal-tracking activity during 8 min non-contingent nicotine-alone tests. Nicotine-evoked goal-tracking was further assessed by calculating the area bound under the nicotine-evoked response curve. The area under the curve (AUC) was calculated from dipper entry duration data during a one-minute interval that was sampled after the initial 15 s (bin 17) following a nicotine infusion. The AUC was calculated relative to a baseline that was derived from the two time bins – one immediately before and one immediately after this one-minute interval (average of bins 17 and 21). This 1 min interval was chosen based on the pattern of responding of sham control rats that was observed after 20 sessions of conditioning phase. Specifically, the dipper entry duration of sham control rats on the final nicotine-alone test (test 4) started to increase from the baseline during the time bins 18 and 19, following by gradual decrease during bins 20 and 21. This gradual increase and decrease in dipper entry duration formed an inverted U-shaped curve with the apex of responding during time bin 19 – 30-45 s after non-contingent nicotine infusion or the time when sucrose is available during conditioning sessions. Responding on bins 17 and 21 forms a baseline of that inverted U-shaped curve. We used this pattern of responding of sham control rats as a guide to forming a nicotine-response curve with the apex at bin 19 and baseline at bins 17 and 21. The baseline was factored into account for individual variation in responding during test sessions. The AUC was calculated using the trapezoid rule (Gagnon and Peterson, 1998; GraphPad Software, 2020; Jaki and Wolfsegger, 2009).

### 2.11 Statistical analyses

Analyses of variance (ANOVA) were used to assess grouped effects. To ensure similar levels of nicotine-sucrose pairings at each of the four tests, only rats that averaged more than five sucrose retrievals per session for the two sessions immediately preceding each test (except for the test one where five sucrose retrievals on a single preceding session were required) were included in the analysis. This approach controlled for the number of total conditioning sessions and for the nicotine exposure among all rats during the study. The relationship between the AUC during test 4 and the size of p-dmCPu lesions was assessed using linear regression with significance determined by the deviation of the calculated slope from zero. Statistical analyses were performed using GraphPad Prism (8.4.3). Significance was declared when *p*-values were less than 0.05.

## 3 Results

### 3.1 Nicotine self-administration: conditioning phase

All rats (sham n=14; a-dmCPu n=11; p-dmCPu n=9) rapidly acquired nicotine self-administration. An omnibus analysis of responding during the self-administration phase revealed no effect of lesion condition on lever presses (p=0.49). Overall, there was a main effect of lever (F_1,76_ = 762.7, *p* < 0.0001), a main effect of session (F_5.7,422.2_ = 5.38, *p* < 0.0001), and significant lever by session interaction (F_20, 1470_ = 15.48, *p* < 0.0001). The difference between active and inactive levers was evident on sessions 2-21 (Bonferroni’s tests; Figure 2A). The retrieval of sucrose did not differ from the number of earned sucrose presentations over the course of the conditioning phase (*p* = 0.26; Figure 2A). On session twelve, a subset of rats (n=5) received the wrong lever assignments. The data from these five rats on session twelve were excluded from the analysis.

**Figure 2.**
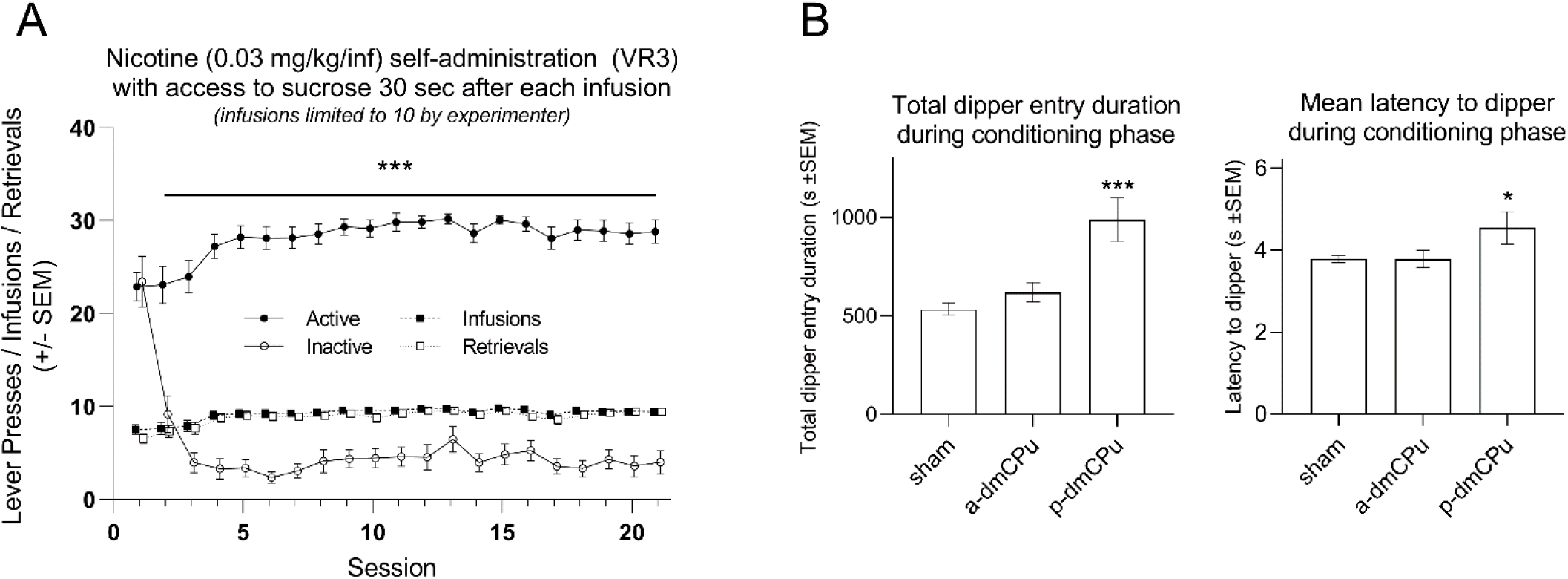
(A) Lever presses, infusions earned, and sucrose retrievals (± SEM) over 21 consecutive self-administration sessions. By the second day of conditioning phase (session 2), rats readily discriminated between the active and inactive lever (*** indicates significant difference; p<0.001). By the fourth day of conditioning phase, all rats averaged ? 9 infusions and sucrose retrievals during each 2-hour self-administration session out of possible 10. (B) The left panel shows the total duration of dipper entries (*** indicates significant difference from sham controls; p<0.001). The right panel shows mean latency to enter the dipper area after nicotine infusions during nicotine self-administration sessions (* indicates significant difference from sham controls; p<0.05).

The analysis of total dipper entry duration throughout the conditioning phase showed a significant main effect of lesion condition (F_2, 742_ = 13.04, *p* < 0.0001). Rats with lesions to p-dmCPu, but not those with lesions to a-dmCPu, showed significantly higher total dipper entry duration throughout the conditioning phase than sham controls (Dunnett’s tests; Figure 2B left panel). In addition, the analysis of mean latency to a dipper after nicotine infusions during the conditioning phase showed a significant effect of lesion condition (F_2, 6941_ = 3.09, *p* < 0.05). Rats with lesions to p-dmCPu, but not those with lesions to a-dmCPu, showed significantly higher mean latency to a dipper after nicotine infusions during the conditioning phase than sham controls (Dunnett’s tests; Figure 2B right panel). These findings suggest that rats with lesions to p-dmCPu had deficits in learning a temporal association between nicotine infusion and a subsequent sucrose delivery resulting in spending more time in the sucrose receptacle than sham controls. These deficits in learning a temporal association between nicotine infusions and sucrose presentations are also supported by increased latency to reach sucrose receptacle after nicotine infusions when compared to sham controls.

### 3.2 Non-contingent nicotine-alone tests: AUC analysis

Left panels of Figure 3 show dipper entry duration by 15 s time bins during the course of the 8 min non-contingent nicotine alone test. Recall that rats were placed into the conditioning chambers, and 1 s nicotine infusion was administered non-contingently to all rats after an initial 4 min of the test (habituation). Rats then remained in the chamber for the rest of the 8 min test (additional 4 min). Throughout the test, the responding in the form of dipper entry duration (snout entry into a receptacle where sucrose was delivered during conditioning sessions; main dependent measure) was recorded by 15 s bins. The main objective during this test was to assess the effect of lesions on nicotine-evoked responding in the form of dipper entry duration (time spent in the sucrose receptacle; goal-tracking) following the nicotine infusion. The AUC analysis was used to assess this nicotine-evoked response following the nicotine infusion. This analysis showed no effect of lesion condition during test 1 (*p* = 0.19; sham n=7, a-dmCPu n=5, p-dmCPu n=5; Figure 3A), test 2 (*p* = 0.68; sham n=9, a-dmCPu n=10, p-dmCPu n=7; Figure 3B), or test 3 (*p* = 0.98; sham n=14, a-dmCPu n=10, p-dmCPu n=7; Figure 3C). However, there was a main effect of lesion condition during test 4 (F_2,31_ = 3.32, *p* = 0.049; Figure 3; sham n=14, a-dmCPu n=11, p-dmCPu n=9). Rats with lesions to p-dmCPu showed significantly lower nicotine-evoked dipper entry durations (significantly smaller AUC) than sham controls (Dunnett’s tests; Figure 3D). The nicotine-evoked goal-tracking of rats with lesions to a-dmCPu did not differ from sham controls on test 4.

**Figure 3.**
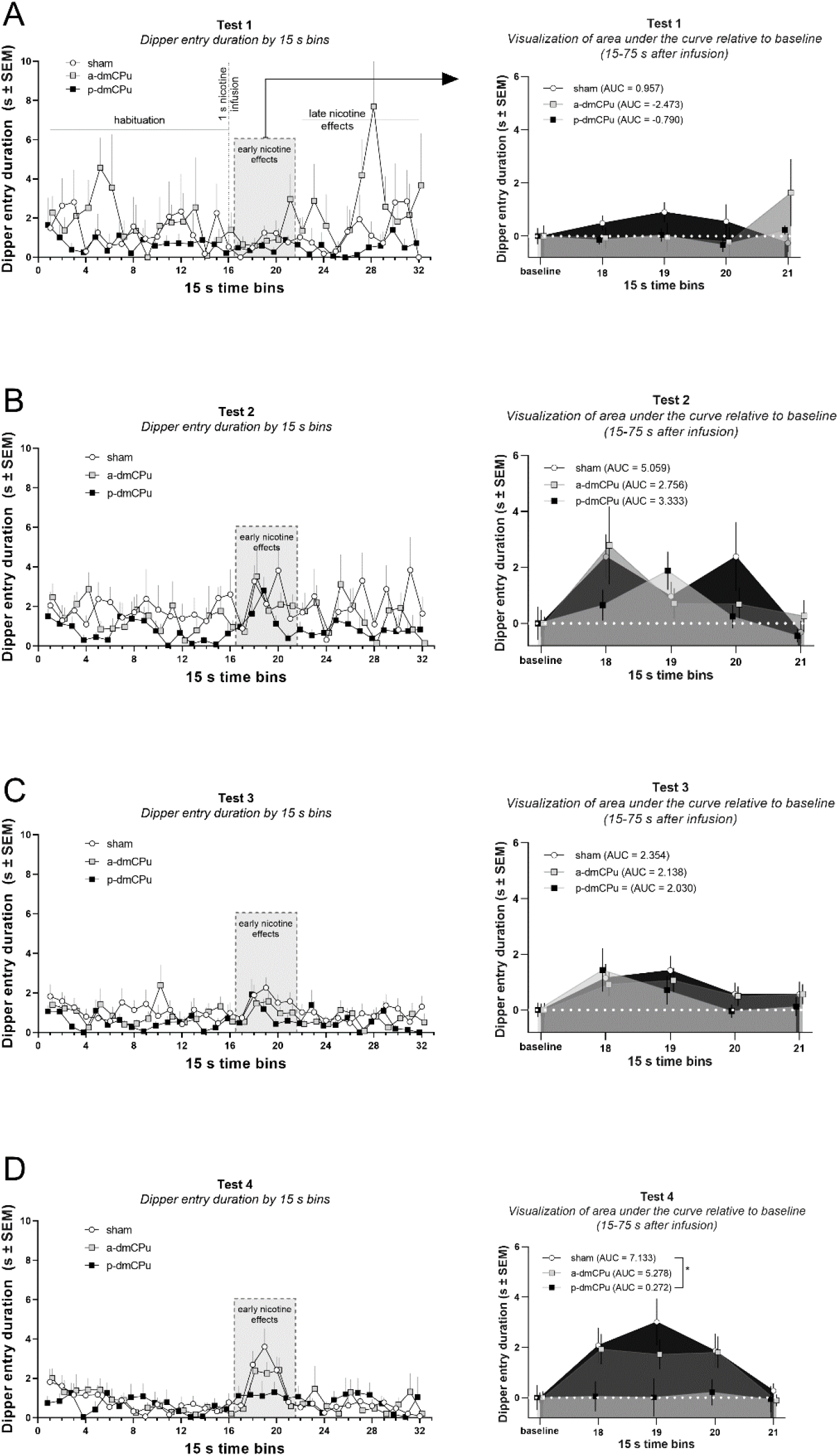
Left panels show dipper entry durations by 15 s bins during non-contingent nicotine-alone tests. The shaded area on left panels outlines goal-tracking on bins 17-21, which represents responding 15-95 s after a sole nicotine infusion administered after 4 minutes of habituation to the testing conditions. Right panels show the calculated area under the curve (AUC; bins 18-21) that was formed by a change in responding from the baseline (dashed white line; * significant difference from responding of sham controls).

To better understand the significant effect of lesions on nicotine-evoked goal-tracking observed during test 4, we conducted a follow-up analysis to assess the role of individual variation in lesion size in the magnitude of observed nicotine-evoked goal-tracking. One of the inherent limitations associated with group analyses presented above is the increased variance introduced by the variation in lesion size. The variance in lesion size is often introduced through uneven dispersion of intracranial lesion infusate that varies in each individual case. As a result, this variation may contribute to variance in observed behavioral measures. With this in mind, we hypothesized that if lesion size significantly contributed to variance observed in nicotine-evoked goal-tracking during test 4, then rats with larger lesions to p-dmCPu will have more pronounced learning deficits in comparison to rats with smaller lesions to the same area. A simple linear regression was used to predict AUC on test 4 based on lesion size (pixels). This assessment showed that rats with larger lesions to the p-dmCPu had more deficits in learning nicotine-sucrose association over the course of conditioning phase (F_1,7_ = 7.346, *p* = 0.03; Figure 4) and that the lesion size explained 51.2 % of the variance (R^2^ = −0.512) in AUC on test 4. These results further confirm that p-dmCPu and not a-dmCPu is involved in the acquisition of associative learning with self-administered intravenous nicotine stimulus. These results also suggest that the deficits in this associative learning as a result of lesions administered to p-dmCPu emerge after a prolonged history of nicotine-sucrose pairings (20 sessions or 200 available nicotine-sucrose pairings) and are dependent on lesion size.

**Figure 4.**
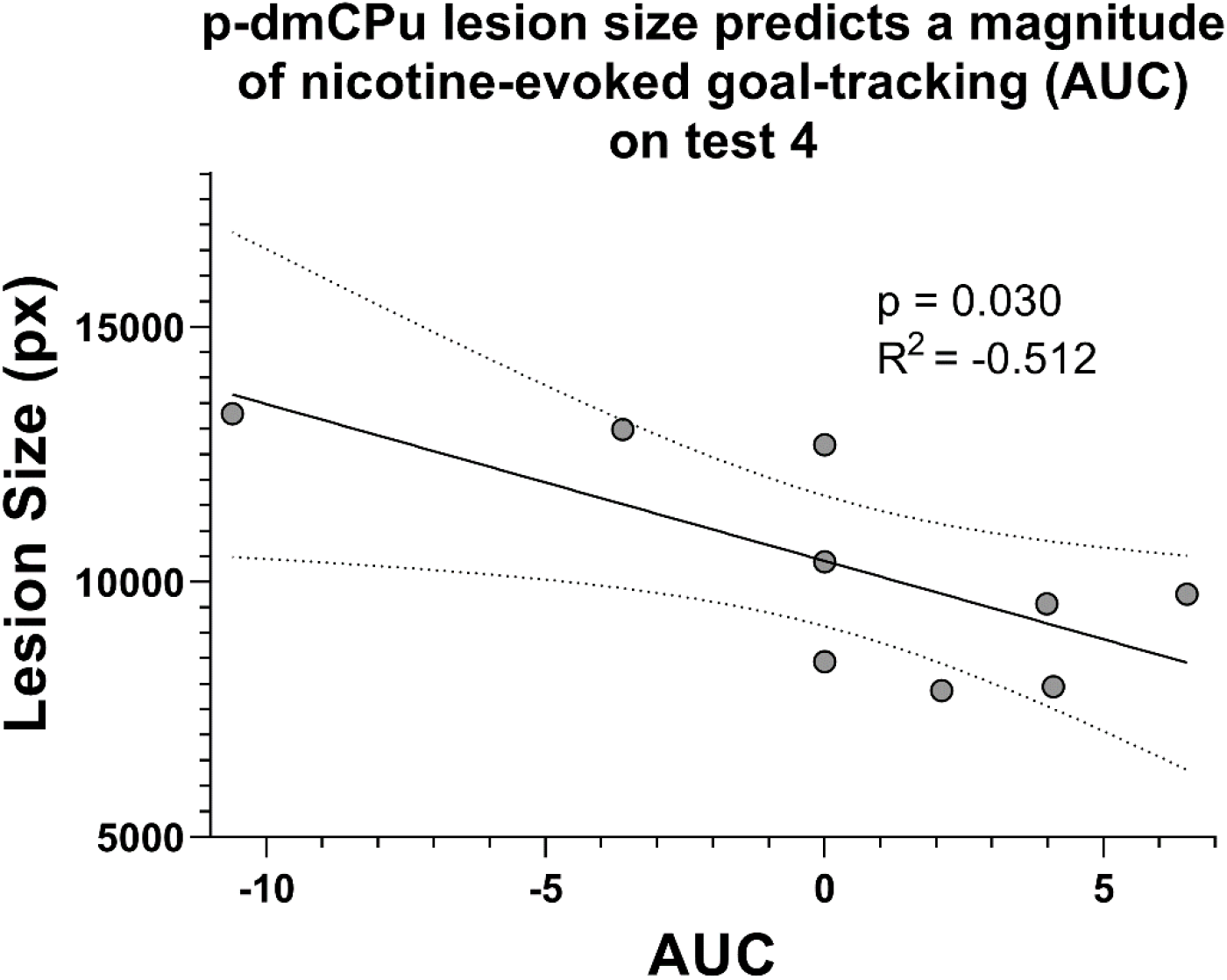
Lesion size explained 51 % of the variance in the area under the curve (AUC; R^2^ = −0.512). Rats with larger lesions having smaller AUC following nicotine infusion on test 4.

### 3.3 Non-contingent nicotine-alone tests: cumulative time course analyses

Within each lesion condition, we divided the nicotine-alone tests into three phases to assess the overall pattern of goal-tracking across all four tests. In addition to providing an assessment of nicotine-evoked goal-tracking emergence over time, this approach also allows us to determine the role of dmCPu in habituation to testing conditions and non-reinforcement because the reward following nicotine infusion is not available during the test. With these goals in mind, we have divided each test session into three distinct phases – habituation (first 4 min before nicotine infusion; bins 1-16), early nicotine effects (1 min after infusion; bins 17-20), and late nicotine effects (last 3 min; bins 21-32). We analyzed dipper entry duration across all tests and phases (4 × 3 mixed-model ANOVA) separately for each lesion condition. Our assessment of this data from rats with sham lesions showed a significant effect of test (F_3,120_ = 4.22, *p* < 0.01), significant effect of phase (F_2,120_ = 3.55, *p* < 0.05), and no significant interaction (*p* = 0.08). Because our main objective was to assess the effects within each phase and test (not assessing any aggregated data like main effects), we followed every ANOVA test with a specific set of a priori post hoc analyses. These post hoc analyses were comparing responding on tests 2-4 to responding on test 1 within each phase and comparing responding during habituation or late nicotine phases to early nicotine effects within each test. Post hoc comparisons of data from rats with sham lesions revealed that goal-tracking on tests 2 and 4 was significantly higher than on test 1 (Tukey HSD tests; Figure 5A; see + symbol). Furthermore, the goal-tracking during early and late nicotine phases on test 4 was lower than goal-tracking during early nicotine during that test (Tukey HSD tests; Figure 5A; see * symbol). Assessment of data from rats with lesion to a-dmCPu revealed a main effect of test (F_3,96_ = 3.49, *p* < 0.05), no effect of phase (*p* = 0.75), and significant interaction (F_6,96_ = 2.66, *p* < 0.05). Post hoc comparisons of that data revealed that goal-tracking during the late phase of the test was lower on tests 3 and 4 when compared to test 1 (Tukey HSD tests; Figure 5B; see + symbol). Furthermore, goal-tracking during the late nicotine phase on tests 1 and 4 was significantly different than goal-tracking during the early nicotine phase on each corresponding test (Tukey HSD tests; Figure 5B; see * symbol). Assessment of data from rats with lesions to p-dmCPu revealed no effect of test (*p* = 0.14), significant effect of phase (F_2,72_ = 3.24, *p* < 0.05), and no interaction (*p* = 0.52). Pre-planned post hoc comparisons revealed that goal-tracking during the early nicotine phase was higher during test 2 than during test 1 (Tukey HSD tests; Figure 5C; see + symbol). Furthermore, the goal-tracking during the habituation phase on tests 2 was significantly lower than goal-tracking during the early nicotine phase during that test (Tukey HSD tests; Figure 5B; see * symbol).

**Figure 5.**
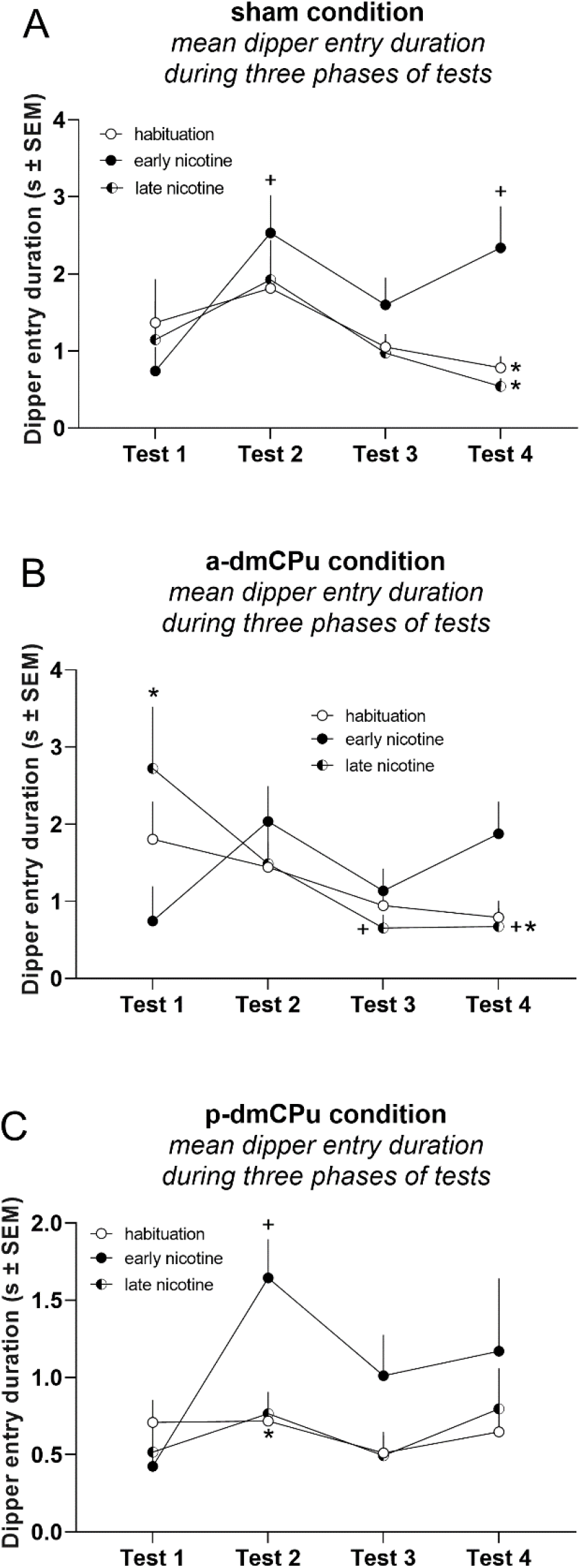
Analysis of dipper entry duration during 8-minute nicotine alone tests across three phrases of the test (habituation, early nicotine, late nicotine). + indicates a significant difference in responding from test 1 for each phase. * indicates a significant difference of responding from corresponding early nicotine phase data point for each test.

## 4 Discussion

Our previous studies using a non-contingent nicotine administration in the context of a discriminated goal-tracking task implicated dmCPu in associative learning with nicotine stimulus (Charntikov et al., 2017, 2012). The goal of the current study was to extend our previous findings to a self-administration model of learning with nicotine stimulus where every contingent nicotine infusion is paired with access to sucrose and where learning is assessed over the course of the study. Using this approach, we found that all rats readily acquired nicotine self-administration and learned to retrieve sucrose from a receptacle at equal rates. However, rats with lesions to p-dmCPu demonstrated blunted learning with self-administered nicotine stimulus, as evidenced by their performance on several measures collected throughout the study. Specifically, rats with lesions to p-dmCPu showed a)increased overall total dipper entry duration during the conditioning phase (Figure 2B, left panel), b) increased overall latency to a sucrose receptacle after nicotine infusions during the conditioning phase (Figure 2B, right panel), c) blunted nicotine-evoked goal-tracking in a non-contingent nicotine-alone test after 20 consecutive sessions of nicotine-sucrose pairing (Figure 3D, right panel), d) blunted increase of nicotine-evoked goal-tracking over time (Figure 5C), and e) relatively low context-induced excitation during non-contingent nicotine-alone tests (Figure 5C, compare to 5A). Moreover, the deficits in associative learning with self-administered nicotine stimulus in rats with lesions to p-dmCPu were correlated with lesion size. This correlation indicates that some of the observed variance in behavioral responses throughout the study could be partially explained by the topography of lesion placement and size. Our findings here are consistent with previous studies, in which rats with lesions to p-dmCPu demonstrated deficits in instrumental and operant learning tasks (Corbit et al., 2012; Corbit and Janak, 2010; Hart et al., 2018; Murray et al., 2012; Yin et al., 2005b) and in nicotine discriminated goal-tracking task (Charntikov et al., 2017). For the first time, we demonstrate the involvement of p-dmCPu in associative learning with nicotine stimulus using a paradigm where rats voluntarily self-administer nicotine and where each nicotine infusion is paired with the access to sucrose. Altogether, our findings confirm the role of p-dmCPu in appetitive learning with nicotine stimulus and further extend previous findings to a model that closely resembles behaviors observed in the clinical population.

While nicotine itself is a weak reinforcer (Dougherty et al., 1981; Henningfield and Goldberg, 1983), it can acquire additional conditioned properties when it comes into association with other reinforcers in the environment. These conditioned properties may underlie the tenacity of the nicotine use habit. For example, in the context of discriminated goal-tracking task rats receive nicotine (the CS) paired with intermittent access to sucrose (the US); on intermixed saline days, sucrose is not available. Across sessions, non-contingent nicotine comes to evoke a goal-tracking response (Besheer et al., 2004; Bevins and Palmatier, 2004; Charntikov et al., 2012; Murray and Bevins, 2009). Behaviorally, this learning follows many of the postulates of Pavlovian conditioning and likely simulates elements of associative learning in human smokers. In this study, we used a self-administration paradigm to model nicotine-reward pairings observed in the clinical population. Using this approach, we show that rats can learn nicotine-reward association when nicotine is self-administered and where each infusion is followed by access to sucrose. We show that rats quickly learn lever discrimination, retrieve over 80 % of available rewards, and show a goal-tracking pattern of behavior after a few training sessions. These findings confirm previous reports demonstrating that non-contingent and contingent intravenous nicotine can serve as a conditioned stimulus, acquire control of behavior, and enhance reinforcing effects of nicotine after a period of learning (Charntikov et al., 2020; Murray and Bevins, 2009). In the present study, we further improved our ability to assess learning with nicotine stimulus by implementing non-contingent nicotine-alone tests that eliminate alternative explanations for the observed nicotine-evoked goal-tracking response. Because each non-contingent nicotine-alone test constitutes extinction learning with nicotine stimulus, we limited the number of tests to four and we followed each test by a training session where rats allowed self-administer nicotine paired with access to sucrose. This improved paradigm allows us to measure learning progress over time and allows for functional assessment of neural substrates underlying learning with nicotine stimulus.

The dmCPu is a heterogeneous structure that plays an important role in regulating goal-directed behaviors (Balleine et al., 2007). The role of dmCPu in learning processes has been extensively studied using instrumental tasks and a number of tests including outcome devaluation or contingency degradation (Balleine et al., 2007; Balleine and Dickinson, 1998; Hart et al., 2018; Yin et al., 2004, 2005b). In the outcome devaluation test, a reinforcer like liquid sucrose or food pellets is devalued by providing ad libitum access to the reward before the test. This food satiation before the test usually leads to a lower incentive to pursue that reward during the test. In a contingency degradation test, a contingency of response for a reinforcer is degraded by weakening the correlation between food delivery and an instrumental response. For instance, in rats trained to press a lever for liquid sucrose or food pellets, the outcome is degraded by delivering extra non-contingent rewards throughout the test. The delivery of “free” rewards during the test leads to a degradation of association between the lever and the reward (Yin et al., 2005b, 2006). Lesions to both the dmCPu and cortical areas with high connectivity with the dmCPu, such as the prelimbic area of the prefrontal cortex, result in impaired contingency learning where rats become insensitive to outcome devaluation or contingency degradation (Balleine and Dickinson, 1998; Yin et al., 2008). Conversely, lesions to the dorsolateral CPu (dlCPu) preserve outcome expectancy while disrupting the formation of habitual behaviors (Yin et al., 2006, 2004). Similarly, rats with transient lesions to the dmCPu demonstrated insensitivity to alcohol devaluation after two weeks of alcohol self-administration, whereas lesions to the dlCPu increased devaluation degradation after eight weeks of alcohol self-administration (Corbit et al., 2012). These reports show that dmCPu plays an important role in a variety of habitual and goal-directed behaviors, including modeled drug-taking (Balleine and O’Doherty, 2010).

Within the dmCPu, there is a further functional subdivision along the anteroposterior axis. The a-dmCPu appears to be important for cue discrimination and cognitive flexibility. For instance, tetracaine hydrochloride inactivation of the a-dmCPu impaired learning in a plus-maze task in which rats were tasked with learning a rule shift between body orientation and visual cue-based response to obtain a food reward (Ragozzino et al., 2002). This effect appears to be mediated by dopamine transmission, as 60OHDA lesions to the a-dmCPu impaired reversal learning where one reward contingency was degraded while another was amplified (Grospe et al., 2018). Conversely, lesions to the p-dmCPu abolish sensitivity to both reward devaluation and outcome contingency degradation, while lesions to the a-dmCPu do not (Yin et al., 2004). NMDA receptor inactivation in the p-dmCPu similarly inhibits reward devaluation, but only when administered prior to a training session rather than prior to the devaluation tests, suggesting that the p-dmCPu is important for action-outcome learning (Yin et al., 2005a). Indeed, rats with transient lesions to the p-dmCPu expressed deficits in both instrumental and operant learning tasks where auditory cues or lever responses were devalued (Corbit and Janak, 2010). Thus, the a-dmCPu is integral for the expression of reward associations, whereas the p-dmCPu is integral for the acquisition of those associations. In our study, after twenty nicotine self-administration sessions, we did not observe any behavioral deficits in rats with lesions to the a-dmCPu (Figure 3D, gray squares). This outcome from our study suggests that a-dmCPu is not involved in appetitive learning with nicotine stimulus. Conversely, rats with lesions to the p-dmCPu (Figure 3D, black squares) showed a decrease in nicotine-evoked goal-tracking after twenty training sessions. This is consistent with the deficits seen in previous studies demonstrating that the inactivation of p-dmCPu impairs instrumental learning (Balleine et al., 2007; Bradfield et al., 2018; Dezfouli et al., 2014; Yin et al., 2005b) and impairs acquisition of learning with nicotine stimulus using the discriminated goal-tracking task (Charntikov et al., 2017). Therefore, our findings further strengthen the understanding that p-dmCPu is central for a broad range of associative learning mechanisms that include learning involving pharmacological states and appetitive stimuli.

To better assess the overall pattern of goal-tracking responding during the non-contingent nicotine-alone tests, we divided the test into three distinct phases for our statistical analyses. These three phases were 1) habituation — the first four minutes of the test during which the rats were nicotine-free; 2) early nicotine effects — the first minute after the nicotine infusion; and 3) late nicotine effects — the remaining three minutes of the test. The assessment of responding during the habituation phase revealed that rats with sham and a-dmCPu lesions decreased their goal-tracking over the four tests. The initial pattern of elevated goal-tracking during the habituation phase is likely evoked by the context that acquired excitatory effects during early sessions of this study. Recall that during our conditioning sessions, the chamber is always paired with nicotine stimulus and likely acquires excitatory properties through the associative learning mechanisms (Bouton and Nelson, 1998; Nelson, 2002). Over the course of four tests, this excitatory response is likely weakened through the nonreinforced (extinction) learning because sucrose is not delivered during the 8-min test. It is also possible that the decrease in this context excitation during tests can be explained by the fact that nicotine stimulus progressively gains the association with sucrose reinforcement and becomes a more salient stimulus than a chamber in triggering a goal-tracking response. The increase in nicotine-evoked goal-tracking (early nicotine effects) is evident by test 2 in rats with sham lesions, and by the final test 4 only rats with sham and a-dmCPu lesions show higher goal-tracking compared to test 1. The responding in the last 3 minutes of the test also gradually decreased over time, suggesting learning about non-reinforcement following a sucrose delivery during conditioning sessions (8-min timeout). Importantly, the half-life of nicotine and its major metabolites are between 97 and 449 minutes in rats (Craig et al., 2014). Therefore, it is unlikely that the observed decrease in responding during the last phase of the test is due to the decrease in nicotine blood concentration. Overall, the assessment of responding in the distinct phases of the non-contingent nicotine-alone test provides a deeper understanding of behavioral responses to contextual cues and to interoceptive nicotine stimulus across time.

In the current study, our goal was to determine whether the a- or p-dmCPu differentially involved in the acquisition of learning with nicotine stimulus. Our findings here demonstrate the importance of the p-dmCPu in learning with nicotine stimulus using contingent nicotine administration. Our findings using this self-administration model of appetitive learning with nicotine stimulus confirm the results of our previous study where rats with lesions to p-dmCPu showed blunted acquisition of learning with non-contingent nicotine stimulus in a context of discriminate goal-tracking task (Charntikov et al., 2020). The dmCPu receives excitatory input from the cortex, and some cortical areas like the prelimbic (PL) region have been implicated in the acquisition of goal-directed actions (Balleine and Dickinson, 1998; Balleine and O’Doherty, 2010; Corbit and Balleine, 2003; Ostlund and Balleine, 2005). It is likely that cortical projections from PL area of the cortex to p-dmCPu are also involved in learning with nicotine stimulus. For example, a bilateral disconnection of PL from the p-dmCPu impedes the acquisition of goal-directed actions (Hart et al., 2018). In that study, rats were first trained to lever press for two different food reinforcers on two different levers and then were assessed for goal-directed actions using a devaluation test where all rats were prefed on one or the other outcomes earned during the training. In that study, rats with both ipsilateral and contralateral disconnections of PL to p-dmCPu showed significant impairment in goal-directed action during the devaluation test (Hart et al., 2018). With this in mind, previous studies provide evidence that the PL→p-dmCPu pathway may be critically involved in learning with pharmacological stimuli. Future studies will need to assess the role of the corticostriatal pathway in appetitive learning with nicotine stimulus to better understand mechanisms underlying nicotine use.

## 5 Acknowledgments

This research and S. Charntikov was supported by NIGMS (GM113131).

